# No evidence of reactive avoidance of baboons (*Papio ursinus* and *Papio anubis*) to the presence of predators

**DOI:** 10.1101/2025.04.01.646592

**Authors:** N. van Rooyen, L. Thel, J. A. Venter, H. Fritz, F. Prugnolle, V. Rougeron

**Author notes:** Corresponding authors: Rougeron Virginie, International Research Laboratory, REHABS, CNRS-NMU-UCBL, Sustainability Research Unit, George Campus, Nelson Mandela, University, George, South Africa,<u>/</u>; Nicholas van Rooyen, Department of conservation management, faculty of science, Nelson Mandela University George Campus, Nelson Mandela University, George, South Africa. Co-first authors: N. van Rooyen, L. Thel. Co-supervised the work: F. Prugnolle, V. Rougeron.

## Abstract

Predators exert strong selective pressure on prey species, shaping their behavioral adaptations. Prey species use proactive responses, such as site selection and the adjustment of daily activity patterns to anticipate and avoid predation exposure. In contrast, reactive responses, including fleeing, referential signaling and retaliation, occur after encountering a predator to mitigate the immediate predation risk. The presence of a predator in an area can also generate longer-term responses, such as reactive avoidance, defined here as the temporary avoidance of the area following the passage of a predator. Such a long-term reactive response remains understudied in primates. To investigate reactive avoidance, we analyzed an extensive camera trap data set of 3,042 detections (defined here as a motion-triggered event of three pictures per trigger during the day and one picture at night) of both chacma (*Papio ursinus*) and olive (*Papio anubis*) baboons, including 6,958 detections of their main predators, lions (*Panthera leo*), leopards (*Panthera pardus*) and spotted hyenas (*Crocuta crocuta*), in four savanna sites (Madikwe Game Reserve, Tswalu Kalahari Private Wildlife Reserve and the Associated Private Nature Reserves, South Africa; Serengeti National Park, Tanzania). We examined whether baboons display species-specific reactive avoidance towards predators up to 72 hours after the detection of a predator using randomization tests. We found no significant decrease in baboon presence (defined as the number of baboon detections) 0 to 24 hours, 24 to 48 hours and 48 to 72 hours after the detection of any predator species. These results suggest that baboons may not display reactive avoidance and rely on alternative predator-avoidance strategies, such as proactive avoidance or aggressive retaliation, to mitigate predation risk.

## Introduction

Predation exerts a strong selective pressure on prey species due to its impact on individual fitness and survival (Humphries & Driver, 1967; Harvell, 1990; Krebs & Davies, 2009). Prey species have evolved a diverse range of behavioral responses to enhance their survival as they encounter fluctuating predation risk in their environment (Lima & Dill, 1990; Kie, 1999). Behavioural adaptations are inherently flexible and allow prey to modify the nature and intensity of their responses based on their spatial and temporal perception of risk in the surrounding environment (i.e., the landscape of fear, Laundré et al., 2001; Palmer et al., 2022), and can be classified into proactive (Broekhuis et al., 2013; Creel, 2018) and reactive responses (Broekhuis et al., 2013; Creel, 2018; Say-Sallaz et al., 2023).

Proactive responses are behavioral adjustments made in anticipation of predation risk, based on prior knowledge of predator locations and preferred habitats (Creel & Christianson, 2008; Creel et al., 2014). These adjustments, which result in the interaction with resource and refuge availability in the landscape (Willems & Russell, 2009; Suscke et al., 2021), operate over broad spatial and temporal scales such as home ranges and seasonal cycles (Droge et al., 2017). For example, adult female Sumatran orangutan (*Pongo pygmaeus abelii*) with infants and adolescents proactively select nest sites further away from the last visited food tree to reduce the predictability of their movements, lowering the chances of encountering predators who may visit food sites regularly (Sugardjito, 1983). Hamadryas baboons (*Papio hamadryas*) make use of cliff faces (Schreier & Swedell, 2012) as sleeping and refuge sites to limit predator accessibility to the group and increase predator detectability (Anderson, 1998; Bidner et al., 2018). Chacma baboons (*Papio ursinus*) use refuge areas on the edge of urban environments, spending less than 1% of their time within the urban area, as a strategy to avoid human presence (Fehlmann et al., 2017). They also increase vigilance when away from roosting sites or when isolated from group members, enhancing threat detection in vulnerable situations (Fischer et al., 2001). However, when foraging, the item size and complexity of the foraging task might reduce the level of vigilance of the individual (e.g., reduced vigilance when processing small food items requiring more attention, Allan et al., 2024).

Reactive responses, in contrast, are immediate reactions triggered by the detection of a predator (Broekhuis et al., 2013). These responses occur on smaller spatial and temporal scales such as foraging sites and across diel activity periods (Droge et al., 2017), driven by real-time threats. As a short term reactive response, white-faced capuchins (*Cebus capucinus*, Fichtel et al., 2005), vervet monkeys (*Chlorocebus pygerythrus*, Seyfarth et al., 1980) and spectral tarsiers (*Tarsius tarsier*, Gursky, 2007) commonly use referential signaling to alert other group members after the detection of a predator. Bonobos (*Pan paniscus*, Druelle et al., 2020), colobine monkeys (*Colobus guereza*, Von Hippel, 1998) and baboons (*Papio* spp., Cowlishaw, 1994; Isbell et al., 2018) seek refuge in elevated areas (e.g., tree canopies and cliffs) to reduce their vulnerability to terrestrial predators. Chacma baboons also demonstrate aggressive predator mobbing when facing a predator, where the group collectively confronts and harasses the predator to deter the attack (Cowlishaw, 1994; Jooste et al., 2013). In addition to these short-term responses, some taxa also display long-term reactive avoidance, leaving the disturbed area for several hours to several days, as frequently observed in ungulates (e.g., in Burchell’s zebra (*Equus quagga*), Courbin et al., 2016; in red deer (*Cervus elaphus*), Chassagneux et al., 2020). However, it remains unclear whether primates actively move away from sites visited recently by predators for extended periods of time (Allan et al., 2024; Hammond et al., 2025).

Baboons present an ideal study species to explore primate anti-predator behavior. Due to their broad geographic distribution, baboons encounter a diverse range of predators in their habitat such as lions (*Panthera leo*), spotted hyenas (*Crocuta crocuta*), leopards (*Panthera pardus*), cheetahs (*Acinonyx jubatus*), and wild dogs (*Lycaon pictus*) (Busse, 1980; Cowlishaw, 1994; Jooste et al., 2013). Although cheetahs (Hayward et al., 2006a) and wild dogs (Hayward et al., 2006b) avoid baboons, lions, leopards, and spotted hyenas are identified as the primary predators of baboons, contributing significantly to baboon mortality (Busse, 1980; Cowlishaw, 1994), even if baboons are usually not the main prey for these predators (Busse, 1980; Jooste et al., 2013; Isbell et al., 2018).

Leopards and lions are both ambush predators, lions hunting baboons during the day, whereas leopards hunt during both diurnal and nocturnal periods, often targeting baboons at their roosting sites at night (Busse, 1980; Bidner et al., 2018; Isbell et al., 2018). Spotted hyenas are cursorial predators (Trinkel, 2010; Wentworth et al., 2011), which mostly target baboons when the opportunity arises as opposed to actively hunting them (Cowlishaw, 1994; Trinkel, 2010; Wentworth et al., 2011). These different predation strategies are known to trigger different anti-predator responses in ungulate prey species (Berger, 1979; Palmer et al., 2021). Similarly in primates, species-specific responses have been documented in some taxa: spectral tarsiers (Gursky, 2007) and vervet monkeys (Seyfarth et al., 1980) emit different alarm vocalizations to differentiate between aerial and terrestrial predators, male olive baboons (*Papio anubis*) tend to position themselves at the periphery of the group to protect more vulnerable group members such as females and juveniles from cursorial predators (Suire et al., 2023).

So far, most studies on baboon-predator interactions have relied primarily on direct field observations (e.g., Busse, 1980; Hill & Weingrill, 2007; Allan et al., 2021). While these methods offer detailed insights into short-term reactive responses, they may influence animal behavior due to the presence of human observers (predator shelter hypothesis, Shannon et al., 2014). Additionally, they are often difficult to implement over extended periods or under specific conditions such as nighttime, limiting their usability to explore long-term reactive responses (Nowak et al., 2014; Dill & Frid, 2020; LaBarge et al., 2022; Allan et al., 2024). In contrast, camera traps are a non-invasive, uninterrupted monitoring alternative (Newey et al., 2015), which reliably detect large carnivores (Meek et al., 2016) as well as primates (Boyer-Ontl & Pruetz, 2014; Bersacola et al., 2022). Recent studies have largely used camera traps to study predator-prey interactions (Smith et al., 2020), particularly space use (e.g., Cusack et al., 2017; Keim et al., 2019) and activity pattern overlaps (e.g., Linkie & Ridout, 2011; Foster et al., 2013). The widespread adoption of camera traps in recent decades has led to the development of long-term monitoring programs across extensive areas (Swanson et al., 2015; Blount et al., 2021), providing a new source of data for the study of predator-prey interactions (e.g., Hammond et al., 2025). Although promising, are these non-targeted, large-scale data sets adequate for examining long-term primate reactive responses to predators?

In this study, we took advantage of the largest camera trap-based monitoring project in Africa, Snapshot Safari (Swanson et al., 2015; Pardo et al., 2021) and analyzed 10,000 camera trap detections from three sites in South Africa and one site in Tanzania to test whether baboons exhibit species-specific reactive avoidance in response to the presence of predators. We tested three hypotheses related to the interaction between baboons and the characteristics of their three main predators: H1) we expected baboon presence to decrease significantly 0 to 24 hours following the detection of spotted hyena, a terrestrial cursorial predator. Spotted hyenas only opportunistically hunt baboons (Cowlishaw, 1994; Trinkel, 2010; Wentworth et al., 2011) and are highly mobile across the landscape (Kolowski et al., 2007). Baboons predominantly forage in proximity to trees or cliffs (Schreier & Swedell, 2012; Fehlmann et al., 2017) that provide potential refuges in the event of a predation attempt. We predict that the occurrence of spotted hyena should have a weak influence on baboons, with baboons returning to the disturbed area shortly after the detection. H2) we expected baboon presence to decrease significantly 0 to 48 hours following the detection of leopard, a semi-arboreal ambush predator. Leopards can pursue baboons into trees, which may prompt the entire group to relocate for a longer period (Trinkel, 2010; Wentworth et al., 2011). However, leopards are also highly mobile and may not remain in the same area for long periods (Martins & Harris, 2013; Hubel et al., 2018). We predict that leopard presence should affect baboons more strongly than spotted hyena presence, leading to their displacement from the disturbed area with a gradual return following the detection of leopard. H3) we expected baboon presence to decrease significantly 0 to 72 hours following the detection of lion, a terrestrial ambush predator. Although baboons can evade lion predation by retreating to elevated refuges, lions can exhibit limited daily displacement and may remain in the same area for extended periods (Elliot et al., 2014), potentially forcing baboons to relocate for a prolonged duration to maintain their normal activity patterns (particularly for foraging) in safer conditions. We predict that lion presence should most strongly affect baboons, resulting in the longest avoidance period compared to other predators, with a slow return following the detection of lion.

## Methods

### Study site selection and description

In this study, we used data from the camera trap monitoring project *Snapshot Safari*, which spans multiple sites and countries (Tanzania, data publicly available at Dryad: http://dx.doi.org/10.5061/dryad.5pt92, Swanson et al., 2015; South Africa, Pardo et al., 2021). Of the 32 sites surveyed, we included sites where all three target predators were detected along with baboons. The South African study sites included Madikwe Game Reserve (MAD, 24°49’S-26°13’E; Figure 1A), Tswalu Kalahari Private Wildlife Reserve (TSW, 27°14’S-22°23’E; Figure 1B), and the Associated Private Nature Reserves (APN, 24°20’S-31°20’E; Figure 1C), inhabited by chacma baboons (Stone et al., 2013). The Tanzanian study site was Serengeti National Park (SER, 2°19’S-34°49’E; Figure 1D), occupied by olive baboons (Zinner et al., 2011). All four study sites were located within the savanna biome, characterized by open grasslands interspersed with isolated trees (Sankaran et al., 2005).

**Figure 1.**
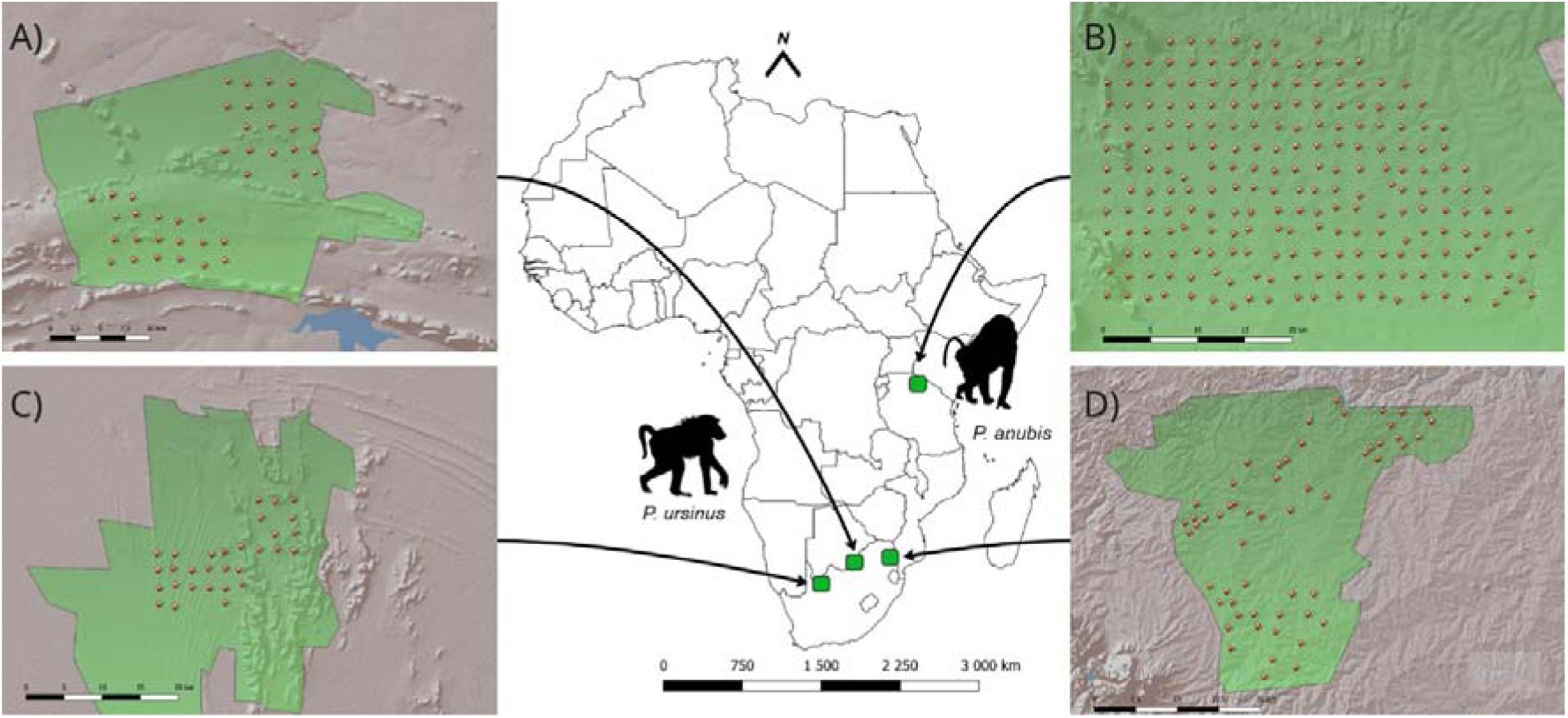
Map of the four study sites in South Africa (data collected between 2017 and 2019) and Tanzania (data collected between 2011 and 2013): A) Madikwe Private Game Reserve (MAD; n = 40 camera traps); B) Serengeti National Park (SER; n = 219 camera traps); C) Tswalu Kalahari Game Reserve (TSW; n = 31 camera traps); D) Associated Private Nature Reserves (APN; n = 56 camera traps). The green boundaries represent the borders of the study areas, the red circles represent the camera trap locations, with their assigned code printed above. Chacma baboon (*Papio ursinus*) occurs in the three South African sites, olive baboon (*Papio anubis*) occurs in the Tanzanian site.

MAD is a 750 km² fenced game reserve located in the Northwest Province, on the border of South Africa and Botswana, within the central bushveld bioregion (Szott et al., 2020). The landscape is characterized by ridges and thorny woodlands dominated by acacia species (Szott et al., 2020). TSW, located in the Northern Cape Province, is a 1,020 km² fenced reserve in the Eastern Kalahari bioregion, with portions extending into the Kalahari dunefields. Elevation ranges from 1,020 m a.s.l. in lower-lying areas to 1,586 m a.s.l. at the highest point (Davis et al., 2010). The vegetation consists of discontinuous annual and perennial grasses interspersed with deep-rooted trees, such as the camel thorn (*Vachellia erioloba*, Beaumont & Bednarik, 2015). APN is a collective of five interconnected reserves (Timbavati, Thornybush, Klaserie, Balule, and Umbabat Private Nature Reserves) covering 1,800 km² in the mopaneveld bioregion of Limpopo and Mpumalanga provinces. It is open to the Kruger National Park (Bedetti et al., 2020). The APN vegetation is highly variable, with isolated trees in a continuous grass understory interspersed with patches of dense woody vegetation (Scholes & Archer, 1997). The SER study site corresponds to an open sub-section of 1,125 km^2^ inside the core of Serengeti National Park, Tanzania, part of the East African savanna biome. It is covered by grasslands interspersed with scattered trees, with steeper slopes to the North-West (Swanson et al., 2015).

### Camera trap data

Data for MAD, TSW, and APN were collected between June 2017 and November 2019 (29 months), and data for SER were collected between June 2010 and May 2013 (35 months). As is common in large-scale monitoring programs, camera traps (n = 357) from different brands (Cuddeback’s Professional Series x-change, white flash in South Africa, and Scoutguard (SG565) incandescent cameras in SER) were deployed. Camera trap placement followed a systematic layout in MAD, TSW, and SER, where camera traps were positioned at the center of a 5 km² grid pattern. Camera trap placement in the APN followed a random allocation layout (*sensu* Meek et al., 2014), based on habitat and terrain structure. The home range size of chacma and olive baboons varies between 24 and 40 km^2^ (Harvey & Clutton-Brock, 1981; Hoffman & O’Rian, 2012; Musyoki & Strum, 2016; Slater et al., 2018). Given that camera traps were deployed at an approximate density of one camera trap per 5 km², the configuration provides good coverage of the baboon groups estimated home range area. We removed five camera traps from the MAD data set that were positioned specifically near waterholes, as predator-prey interactions may likely differ at waterholes due to the trade-off between the need for scarce resources (i.e., water) and predator avoidance (Edwards et al., 2016). In the present study, we used a total of 346 camera traps (n_MAD_ = 40; n_TSW_ = 31; n_APN_ = 56; n_SER_ = 219) for our analyses (see below for data processing and selection steps).

The camera traps were enclosed in steel casings and fixed at a height of approximately 50 cm above the ground (Swanson et al., 2015; Pardo et al., 2021). To minimize glare caused by the rising and setting sun, the camera traps were oriented to face as far North or South as possible. Cameras were also directed based on vegetation cover, avoiding areas of high grass density to reduce the probability of obtaining blank pictures and maximizing animal detection. The sensitivity of each camera trap was set to a medium level (50-75%) to minimize false triggers caused by windblown vegetation. No lures were used to attract any species to the camera trap sites. Each camera trap was programmed to take three pictures for every motion detection during the day and one at night (see Pardo et al., 2021 for a more detailed description of the protocol in the South African sites; and Swanson et al., 2015 for the Tanzanian site). To minimize the impact of potential variability in detection probabilities due to differences between protocols and sites (Cusack et al., 2017; Herrera et al., 2021), we analyzed our four sites separately.

Species were identified in each capture event (i.e., any single picture or consecutive series of pictures initiated by an animal-trigger, regardless of the number of individuals present in the picture(s), Meek et al., 2014) using two methods: (i) for SER, the citizen science program Zooniverse (www.zooniverse.org) which uses classifications from the general public (consensus after 10 matching classifications of species or species combination; see details in Swanson et al., 2015); (ii) for the three South African sites, the online program TrapTagger (www.wildeyeconservation.org), which was used to manually record animal species as identified by trained laboratory technicians. After species classification, we obtained a total of n = 1,625 detections of chacma baboons, n = 1,531 detections of olive baboons, n = 6,929 detections of spotted hyenas, n = 449 detections of leopards and n = 3,886 detections of lions.

As our objective was to evaluate the presence of baboons up to 72 hours after (and symmetrically, before) the detection of a predator, we exclude any predator detection that occurred within the 72 hours following the camera’s deployment and within 72 hours before the end of the camera’s operational lifespan. To ensure consistency, we applied a 30-minute time-to-independence filter to predator detections, retaining the central detection within a cluster of consecutive detections as the main detection for a given predator detection. This threshold aligns with previous studies that showed that increasing the independence interval has minimal impact on observed activity patterns of large carnivores (Linkie & Ridout, 2011; Searle et al., 2021; TjadenLJMcClement et al., 2025). Retaining the central detection instead of the first detection as usually done in the literature ensures symmetry when testing for baboon response to a predator before and after the predator detection in our study design. Our final data set was thus composed of n = 1,584 detections of chacma baboons, n = 1,458 detections of olive baboons, n = 5,071 detections of spotted hyenas, n = 355 detections of leopards and n = 1,532 detections of lions (Table 1).

**Table 1.**
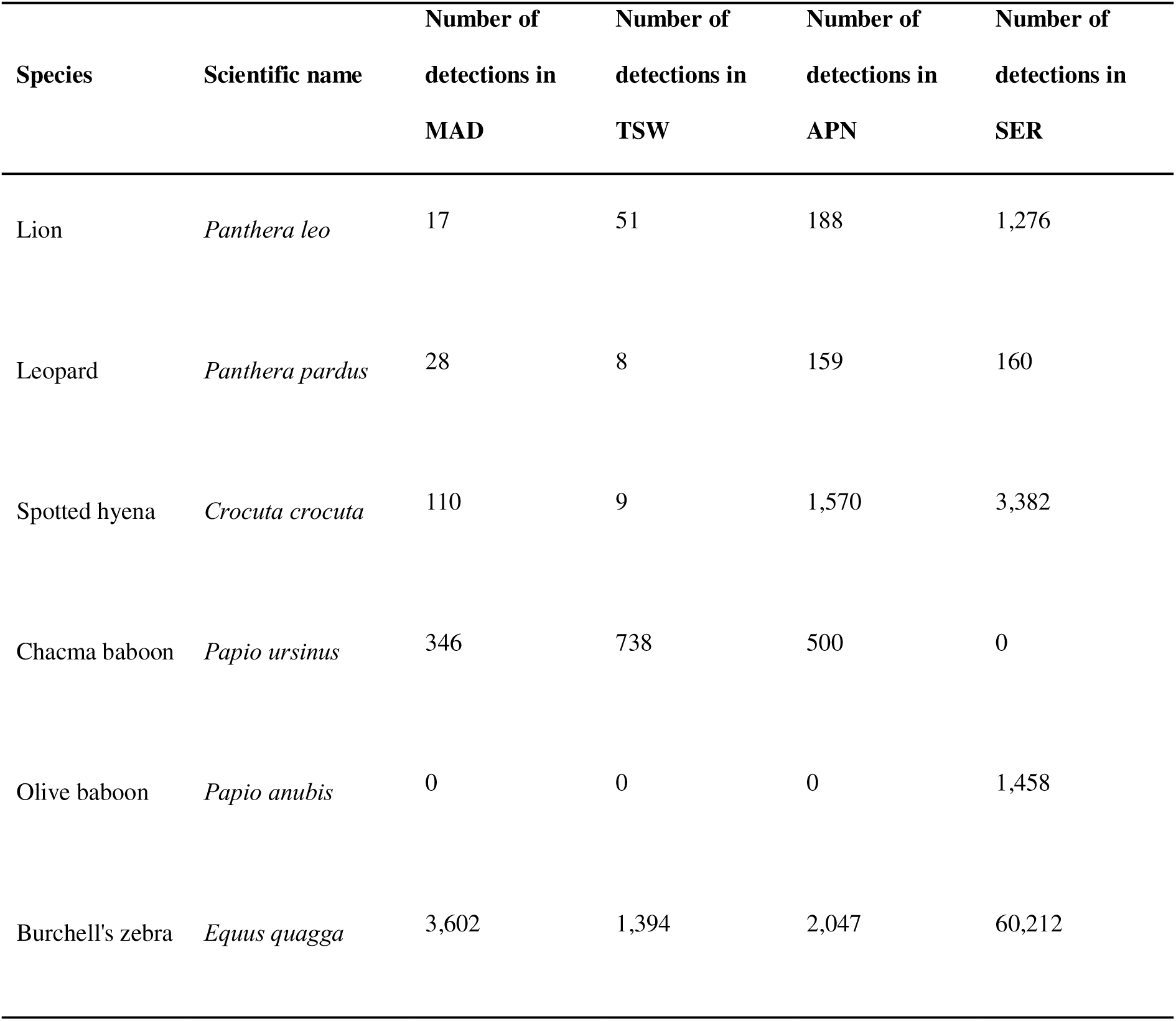
Number of detections for the species of interest: chacma baboon (*Papio ursinus*), olive baboon (*Papio anubis*), Burchell’s zebra (*Equus quagga*), spotted hyena (*Crocuta crocuta*), leopard (*Panthera pardus*), and lion (*Panthera leo*), in Associated Private Nature Reserves (APN), Madikwe Game Reserve (MAD), Tswalu Kalahari Private Wildlife Reserve (TSW) (South Africa, data collected between 2017 and 2019), and Serengeti National Park (SER) (Tanzania, data collected between 2011 and 2013). A 30-minute time-to-independence filter was applied to predator detections (see Methods section for details).

### Statistical analyses

To assess whether baboon presence after a predator detection was lower than expected by chance, we used a randomization test to compare the number of baboon detections after the predator detection to the ones obtained under the assumption of non-avoidance of predators (Niedballa et al., 2019; Zhang & Zhao, 2023). We calculated the total number of baboon detections following predator detections in each site, for each predator species independently. This was done for three consecutive 24-hour time-blocks after predator detection: 0 to 24 hours after the predator detection, 24 to 48 hours after the predator detection and 48 to 72 after the predator detection. Extending the observation period beyond 72 hours after predator detection was avoided due to the difficulty in associating the prey species presence or absence with the original predator detection (Courbin et al., 2016; Swinkels et al., 2023). We further analyzed baboon detections before the predation detection to establish a baseline for their presence, with the expectation of no significant difference in the number of baboon detections. This allowed us to determine the strength of the predator effect while accounting for natural fluctuations in baboon activity. The analysis of baboon detections before a predator detection was also conducted in three 24-hour time-blocks over a 72-hour period.

Predator species have been reported to sometimes follow each other (e.g., kleptoparasitism, Cusack et al., 2017). In our data set, < 5% of the days when a predator was detected at a camera trap site were also characterized by the detection of a second predator species (0.7% in MAD, 0.0% in TSW, 2.5% in APN, 3.6% in SER). We thus considered that there was no significant risk of interpreting baboon reactive avoidance due to a predator species when examining the effect of another species.

To obtain the distribution of baboon detections under the null hypothesis (i.e., non-avoidance of predators), we randomized the dates while preserving the original timing to maintain natural diel activity patterns of each predator detection 1,000 times in each camera trap-roll (i.e., the continuous working time of a camera trap between two services, approximately three months) detecting both baboons and the predator of interest. We then estimated, for each randomized data set, the total number of baboon detections in each time-block. We excluded any camera trap with a lifespan < 30 consecutive days to avoid restricting the randomization process. Predator detections captured by camera traps with a shorter lifespan could only be randomized within a limited time window, which would produce unreliable results.

For each 24-hour time-block, we evaluated whether the observed number of baboon detections deviated significantly from the assumption of non-avoidance of predators. To do so, we calculated the significance value (*p*) for each time-block as the proportion of iterations in our randomization procedure in which the number of baboon detections was inferior (superior, respectively) or equal to the observed number of baboon detections after (before, respectively) the predator detection.

All data processing and analyses were performed by the authors using the R software (version 4.5.1; R Core Development Team, 2025) with the help of the packages *chron* (James and Hornik, 2024), *lubridate* (Grolemund and Wickham, 2011), *stringr* (Wickham, 2025) and *tidyr* (Wickham et al., 2025). The code to analyze and reproduce this study has been deposited in Zenodo and is available online at https://zenodo.org/records/15018760.

### Validation of the randomization approach

To validate our randomization approach, we tested it on Burchell’s zebras (*Equus quagga)*. Burchell’s zebras are known to display reactive avoidance in response to lions for up to 24 hours after an encounter (Thaker et al., 2011; Courbin et al., 2016; Say-Sallaz et al., 2023), therefore we expected to find a significant response in the 0 to 24 hour time-block, after lion detection.

APN had the highest number of lion detections among the South African sites (n_APN_ = 188, n_MAD_ = 17, n_TSW_ = 51; sample sizes after data selection process). Although SER had higher numbers of both lions (n_SER_ = 1,276) and Burchell’s zebras (n_SER_ = 60,212), analysing these data revealed too computationally intensive using our randomization procedure. Additionally, the number of Burchell’s zebras (n_APN_ = 2,047) was of the same magnitude as the number of baboons in SER, which represents our largest data set in terms of baboon detections (n_SER_ = 1,458), making it a suitable comparison point to assess the effect of sample size on the results.

To assess the effect of sample size on the analyses, we performed 100 random subsets of Burchell’s zebra detections in APN to match the number of baboon detections (n_APN_ = 500). We thus created a down sampled data set, for which we conducted the randomization analysis for each of the 100 iterations and calculated the frequency of significant versus non-significant values.

## Results

### Method validation: reactive avoidance in Burchell’s zebras

As expected, Burchell’s zebras showed significant reactive avoidance 0 to 24 hours after lion detection in APN (*p* = 0.024, Figure 2), providing support for the soundness of our randomization approach. We did not find significant reactive avoidance 24 to 48 hours (*p* = 0.176) and 48 to 72 hours (*p* = 0.495) after lion detection (Table 2). After taking a subset of Burchell’s zebra detections to match the number of baboon detections in APN (n = 500), we could no longer detect this response 0 to 24 hours following lion detection (*p (mean ± SD)* = 0.182 ± 0.103, Supplementary Figure 1).

**Figure 2.**
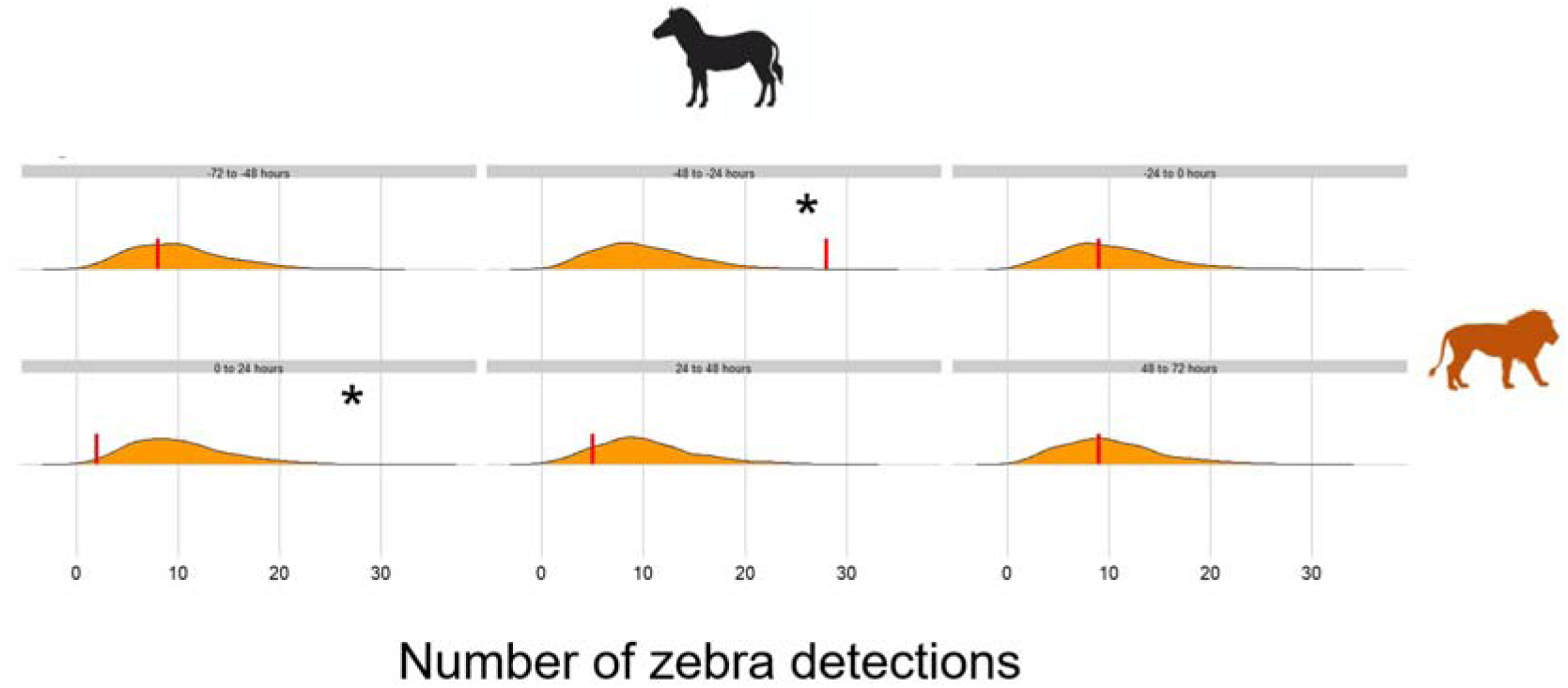
Distribution of Burchell’s zebra (*Equus quagga*) detections obtained from the randomized data sets (n = 1,000) before and after lion (*Panthera leo,* in orange) detection in Associated Private Nature Reserves (APN), South Africa (data collected between 2017 and 2019). Each panel corresponds to one of six consecutive 24-hour time-blocks relative to the predator detection: 72 to 48 hours before, 48 to 24 hours before, 24 to 0 hours before, 0 to 24 hours after, 24 to 48 hours after, 48 to 72 hours after. The red vertical lines represent the observed number of Burchell’s zebra detections for each time-block. The shaded area corresponds to the density of randomization trials, the area under the curve thus being equal to 1. *: significance to the level alpha = 0.05.

**Table 2.** Significance values (*p*) calculated by comparing the observed number of detections of chacma baboon (*Papio ursinus*) and olive baboon (*Papio anubis*) before and after predator detections: spotted hyena (*Crocuta crocuta*), leopard (*Panthera pardus*), and lion (*Panthera leo*), to the distribution of number of detections produced by randomizing predator detections under the assumption of non-avoidance, across six consecutive 24-hour time-blocks relative to the predator detection (72 to 48 hours before, 48 to 24 hours before, 24 to 0 hours before, 0 to 24 hours after, 24 to 48 hours after, 48 to 72 hours after) in the four study sites: Associated Private Nature Reserves (APN), Madikwe Game Reserve (MAD), Tswalu Kalahari Private Wildlife Reserve (TSW) (South Africa, data collected between 2017 and 2019), and Serengeti National Park (SER) (Tanzania, data collected between 2011 and 2013). *: significance to the level alpha = 0.05.

| Study site | Prey species | Predator species | -72 to -48 hours | -48 to -24 hours | -24 to 0 hours | 0 to 24 hours | 24 to 48 hours | 48 to 72 hours |
| --- | --- | --- | --- | --- | --- | --- | --- | --- |
| MAD | Chacma baboon | Lion | 0.503 | 1.000 | 1.000 | 0.900 | 0.469 | 0.452 |
|  |  | Leopard | 1.000 | 0.105 | 1.000 | 0.808 | 0.802 | 0.496 |
|  |  | Spotted hyena | 0.877 | 1.000 | 0.904 | 0.742 | 0.251 | 0.107 |
| TSW | Chacma baboon | Lion | 0.451 | 0.288 | 0.033 * | 0.925 | 0.664 | 0.354 |
|  |  | Leopard | 1.000 | 1.000 | 0.149 | 0.792 | 0.647 | 0.773 |
|  |  | Spotted hyena | 1.000 | 1.000 | 1.000 | 0.912 | 0.714 | 0.722 |
| APN | Chacma baboon | Lion | 0.671 | 0.001 * | 1.000 | 0.526 | 0.251 | 0.373 |
|  |  | Leopard | 0.547 | 0.739 | 0.597 | 0.573 | 0.114 | 0.600 |
|  |  | Spotted hyena | 0.886 | 0.072 | 0.007 * | 0.225 | 0.498 | 0.144 |
| SER | Olive Baboon | Lion | 0.869 | 0.781 | 0.160 | 0.098 | 0.460 | 0.679 |
|  |  | Leopard | 0.551 | 0.434 | 0.012 * | 0.599 | 0.141 | 0.788 |
|  |  | Spotted hyena | 0.574 | 0.317 | 0.008 * | 0.984 | 0.400 | 1.000 |

### Reactive avoidance in baboons

Our results did not show any significant reactive avoidance of baboons in response to the detection of spotted hyenas, across any time-block after predator detection (0 to 24, 24 to 48, or 48 to 72 hours) in any of the three study sites in South Africa (all p > 0.05, Table 2, Figure 3). Similarly, there was no evidence of baboons displaying reactive avoidance following the detection of lions or leopards in any South African site and across any time-block after their detection (all p > 0.05, Table 2, Figure 3). However, in APN, baboon presence increased significantly 48 to 24 hours before the detection of lions (p < 0.001) and 24 to 0 hours before the detection of spotted hyenas (p < 0.007, Table 2, Figure 3). In TSW, baboon presence increased significantly 24 to 0 hours before the detection of lions (p = 0.033, Table 2, Figure 3).

**Figure 3.**
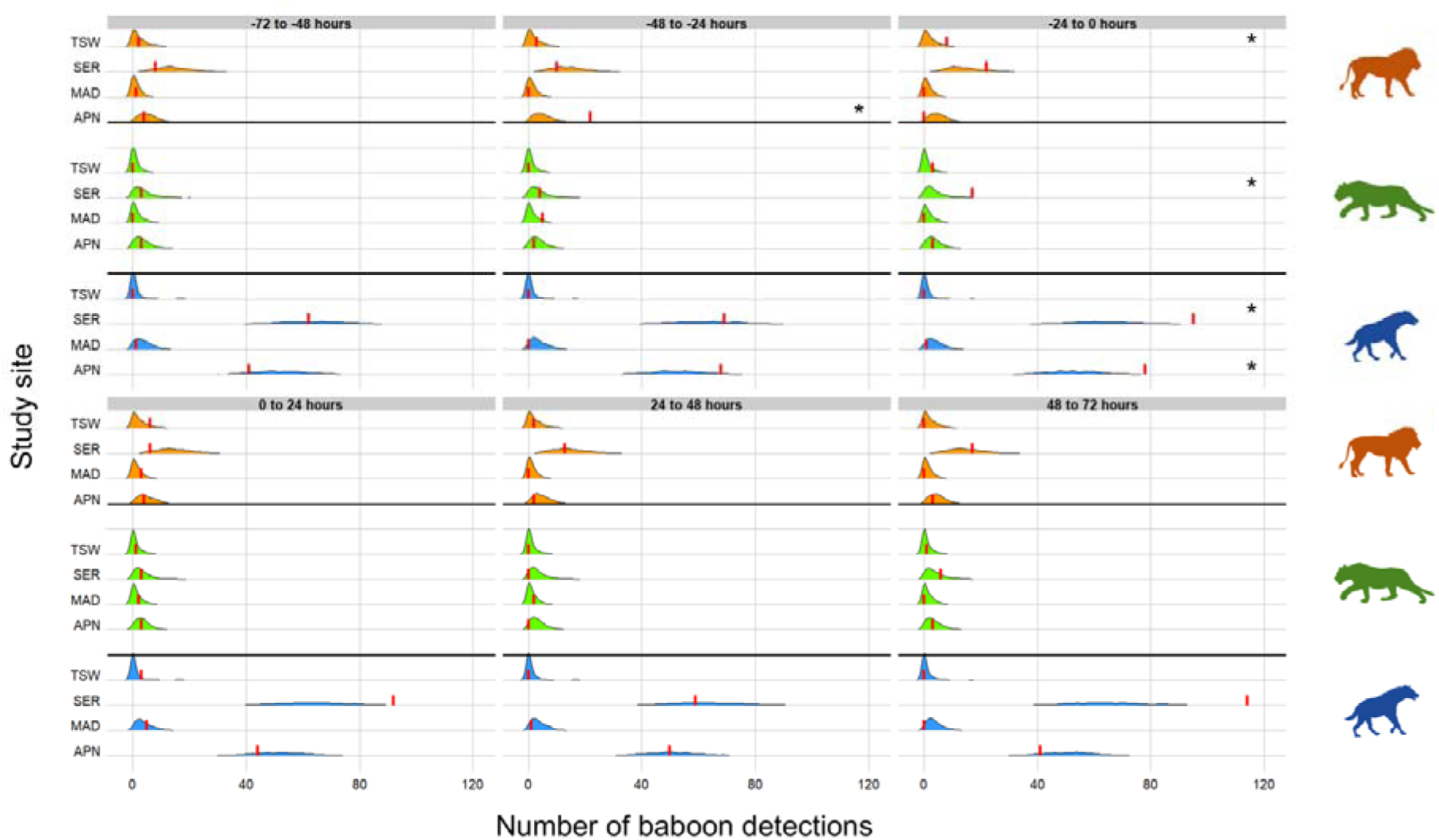
Distribution of baboon (*Papio ursinus* and *Papio anubis*) detections obtained from the randomized data sets (n = 1,000) before and after predator detection: lion (*Panthera leo*, in orange), leopard (*Panther pardus*, in green) and spotted hyena (*Crocuta crocuta*, in blue) across the four study sites: Associated Private Nature Reserves (APN), Madikwe Game Reserve (MAD), Tswalu Kalahari Private Wildlife Reserve (TSW) (South Africa, data collected between 2017 and 2019), and Serengeti National Park (SER) (Tanzania, data collected between 2011 and 2013). Each panel corresponds to one of the six consecutive 24-hour time-blocks relative to the predator detection: 72 to 48 hours before, 48 to 24 hours before, 24 to 0 hours before, 0 to 24 hours after, 24 to 48 hours after, 48 to 72 hours after. The red vertical line represents the observed number of baboon detections for each site/predator/time-block combination. The shaded area corresponds to the density of randomization trials, the area under the curve thus being equal to 1. *: significance to the level alpha = 0.05.

In SER, where the number of baboon detections exceeded those at all South African sites (SER detection rate approximately 4.21 times higher than MAD, 2.92 times higher than APN and 1.98 times higher than TSW, Table 1), and was close to the number of Burchell’s zebra detections in APN, we still did not detect significant reactive avoidance of baboons in response to any predator tested and across all time-blocks after predator detection (all p > 0.05, Table 2, Figure 3). Consistent with observations in APN, baboon detections in SER were significantly higher than expected under the null hypothesis 24 to 0 hours before the detection of spotted hyenas (p < 0.008, Table 2, Figure 3). Additionally, baboon detections in SER were significantly higher than expected under the null hypothesis 24 to 0 hours before the detection of leopards (p = 0.012, Table 2, Figure 3).

## Discussion

Using camera trap data from a large-scale monitoring program in four different savanna sites in South Africa and Tanzania, we found no evidence that either chacma or olive baboons display reactive avoidance for as long as 72 hours after the detection of any of their main predators (lions, leopards and spotted hyenas). Contrary to our expectations, the results showed no difference in the duration of the reactive avoidance response of baboons according to predator characteristics (i.e., predator hunting strategy, predator daily mobility, predator specialization in baboon hunting) at such a time scale, thus providing no support for H1, H2 and H3. Although these results align with the alternative anti-predator strategies commonly used by baboon species, our analyses also suggest a potential lack of statistical power to detect a significant effect due to small sample sizes.

The absence of reactive avoidance at a one-to-three-day timescale suggests that baboons might rely on short-term rather than long-term anti-predator responses. Baboons usually forage in areas with tall trees which can serve as refuge against terrestrial predators such as lions and spotted hyenas (Cowlishaw, 1997). They also exhibit effective sentinel behaviors, one individual monitoring for threats while the rest of the group forages (Horrocks et al., 1986; Bezerra et al., 2008), which leads to the early detection of predation attempts. The danger is rapidly signaled to conspecifics via contact barks (Bailey, 1993; Cowlishaw, 1994; Jooste et al., 2013), which allows for their prompt retreat. Baboons may thus remain in the vicinity of a predator, and retreat into refuges temporarily only in the instance of a predation attempt, rather than completely avoiding a potentially resource-rich area for several days when frequented by a predator. Similarly, Hammond et al. (2025) found no evidence of predator-occupied area avoidance by chacma baboons and suggest that they might instead display short-term reactive avoidance. The modulation of reactive anti-predator responses by environmental variables such as the proximity of refuge areas reducing the vulnerability to predators has also been documented in other species (e.g., in woodchucks (*Marmota monax*), Kramer & Bonenfant, 1997; in European hares (*Lepus europaeus*), Weterings et al., 2016).

Additionally, baboons are notorious for their aggressive retaliation toward predators, particularly leopards (Bailey, 1993; Hayward et al., 2006a: Jooste et al., 2013). A review by Cowlishaw (1994) showed that, when baboons retaliate, they successfully deter predation in 93% of cases, and four out of 11 observed predation attempts resulted in the death of the attacking leopard. This behavior is predominantly exhibited by males in both chacma (Cowlishaw, 1994; Isbell et al., 2018) and olive (MacCormick et al., 2012) baboons, owing to their larger teeth and body size (Virgadamo et al., 1972; Galbany et al., 2015; Johnson, 2003). Similarly, a large number of primate species, mainly males, display mobbing and counter-attack behavior as an anti-predator response (van Schaik et al., 2022). Such a response could decrease predation attempts by lowering the predator’s chance of successful capture and thus, their interest in the species (Lima & Dill, 1990; Alberts, 2018).

Surprisingly, we detected evidence suggesting an increased presence of baboons before a predator detection across all study sites (except MAD). Although these results might be artifacts due to the small sample sizes, they might also be related to the fact that baboons use the ground more frequently in the absence of predators as it is perceived as safer (Hammond et al., 2022; Hammond et al., 2025). This could lead to an increased capture rate of the same individuals during their terrestrial displacements in the monitored area as baboons tend to follow relatively consistent daily foraging routes and regularly revisit the same sites (Noser & Byrne, 2010). Our findings may therefore illustrate a subsequent decline in site use, not because of a decrease in resource availability, but rather due to the recent passage or presence of a predator.

The ability to detect spatio-temporal avoidance confidently when applying a randomization procedure to camera trap data depends on the strength of the avoidance behavior and the number of detections of both species, the latter needing to be as high as 100 detections per species in the case of weak avoidance (Niedballa et al. 2019). In our method validation, we found that Burchell’s zebras showed reactive avoidance toward lions within the 0 to 24 hour time-block after a lion detection, consistent with previous findings in this species (Thaker et al., 2011; Courbin et al., 2016; Say-Sallaz et al., 2023) and thus supporting the validity of our approach. However, after the number of detections of Burchell’s zebras was sub-sampled to match the highest number of baboon detections, we no longer detected reactive avoidance. Sample sizes being relatively low in our four study sites (particularly in MAD and TSW), we might lack statistical power to detect significant reactive avoidance in baboons. To completely rule out this possibility, more camera traps may be necessary to obtain better detection rates of both prey and predators at a given site. Si et al. (2014) and Evans et al. (2019) showed that when studying coyotes, increasing the camera trap density by a single camera at each capture site increases the detection probability by four. Additionally, baboons being capable of utilizing arboreal strata, a mixed design of terrestrial and arboreal camera traps could be adopted to cover all three spatial dimensions of baboon movement and increase their detection rate (Bersacola et al., 2022). Although large-scale camera-trap based monitoring programs present a novel opportunity for preliminary explorations of predator-prey interactions, optimizing camera trap density and placement appears essential to improve detection rates, particularly for low density species with complex habitat use such as primates.

## Conclusion

This study contributes to broadening our understanding of predator-prey interactions in two primate species. The potential absence of reactive avoidance in chacma and olive baboons raises important questions about the variability of their anti-predator strategies, not only in savanna biomes but also across other environments in response to different predators (Zinner et al., 2011; Stone et al., 2013). This study further highlights the influence of detection rates on the detectability of behavioral responses when using camera traps, as illustrated by the loss of significance of reactive avoidance in Burchell’s zebras after the reduction of sample size. These limitations related to camera trap density and placement further emphasize the challenges of accurately quantifying prey-predator interactions among low-density species using systematic and random layouts, common in large-scale monitoring projects.

## Authors’ contributions

Conceptualization: VR, FP – Methodology: NVR, LT, VR, FP – Data acquisition: HF, JV – Formal analysis: NVR, LT, VR, FP – Investigation: NVR, VR, FP, LT – Writing: Original draft: NVR, VR, FP, LT – Writing: Review & Editing: NVR, LT, VR, FP, HF, JV – Supervision: VR, FP – Funding acquisition: VR, JV

## Acknowledgments

The authors thank the CNRS for funding this project. They also thank the Wildlife Ecology Lab, Nelson Mandela University – George campus, CNRS, IRL REHABS, and the Snapshot Safari project for the provision and processing of the data. Funding for Snapshot Safari was provided by the National Research Foundation and the Foundational Biodiversity Information Program (Grant number FBIP170720256205). We would like to thank the Snapshot Serengeti project for providing access to the camera-trap data used in this study, and in particular Craig Packer and the Snapshot Serengeti team for their long-term efforts in data collection and curation. We are also grateful to the many volunteers who contributed to data classification through the Snapshot Serengeti platform.

## Inclusion and diversity statement

The authors are committed to fostering an environment of open dialogue, respect, and cultural inclusion. Our collaboration between African and European researchers reflects our dedication to international scientific exchange and diverse perspectives. Within our research team and laboratory, we actively promote a supportive and inclusive environment. Our first author lives with temporal lobe epilepsy, a condition that affects both memory and language. We acknowledge the unique challenges this presents in research and take meaningful steps to ensure that our team remains understanding and accommodating to this disability. We firmly oppose all forms of discrimination based on sex, gender, race, or any other identity marker. Our partnership is grounded in fairness and merit, and we evaluate all contributions based on the quality and impact of the work, not on the individual’s background or identity.

## Data availability statement

Pictures from the Snapshot South Africa program can be requested at Snapshot Safari upon reasonable request. Pictures from the Snapshot Serengeti program are released under the Community Data License Agreement (permissive variant) and are available from the Labelled Image Library of Alexandria – Biology and Conservation: http://lila.science/datasets/snapshot-serengeti. All classification data and metadata are publicly available at Dryad: http://dx.doi.org/10.5061/dryad.5pt92.

## Code availability statement

The code to analyze and reproduce this study has been deposited in Zenodo and is available online at https://zenodo.org/records/15018760.

## Ethical approval

Ethics approval was not required for this study according to local legislation [Parks and Wildlife Act].

## Consent to participate

Not applicable.

## Consent for publication

Not applicable.

---

**Supplementary Figure 1.**
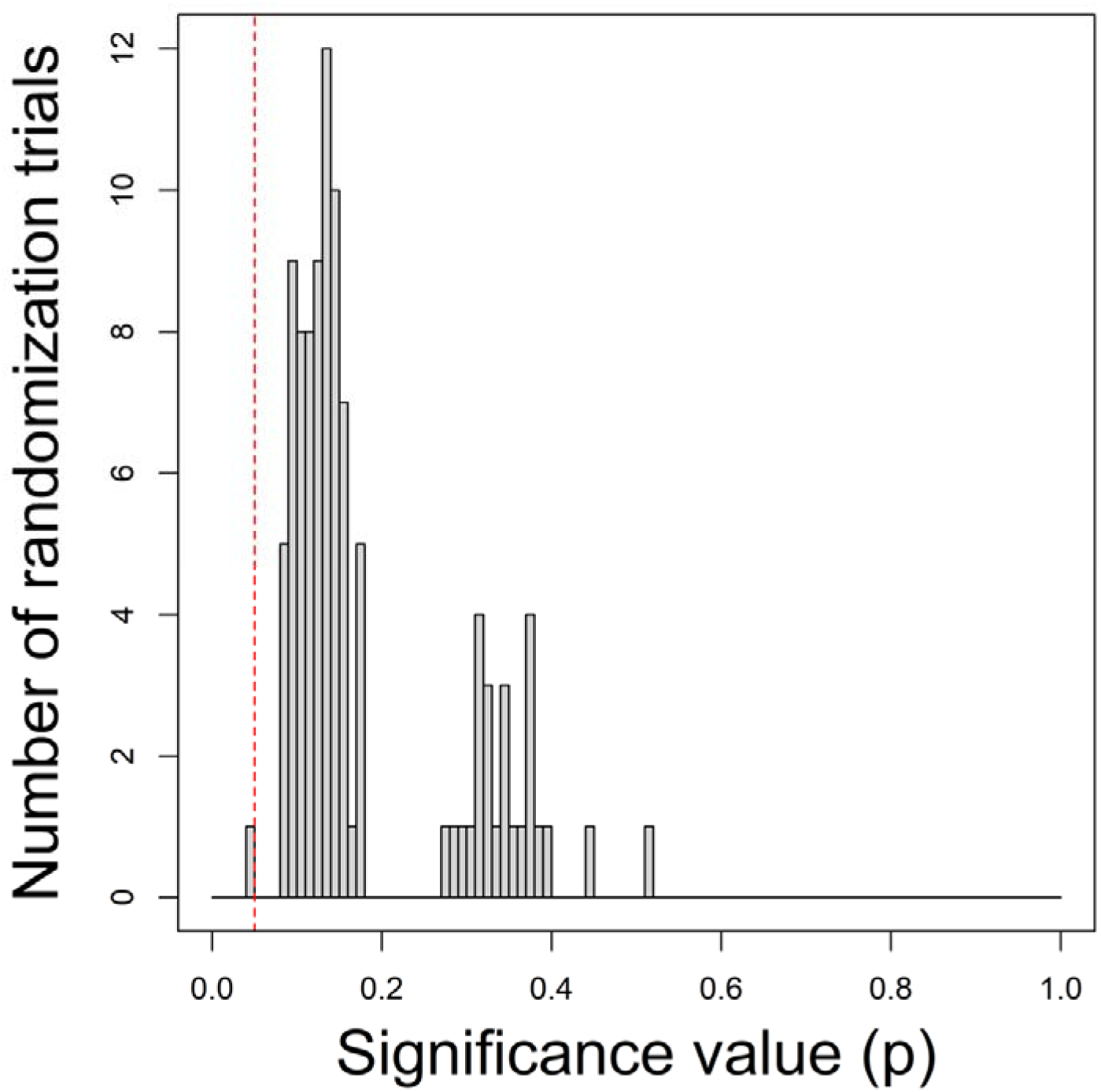
Distribution of significance values (*p*) from the randomization test assessing the effect of lions (*Panthera leo*) on Burchell’s zebra (*Equus quagga*) detections in Associated Private Nature Reserves (APN), South Africa (data collected between 2017 and 2019) within the 0 to 24 hours following lion detection. The test was conducted on randomly down-sampled Burchell’s zebra detections data sets (n = 100). The red dashed line indicates the significance value threshold of alpha = 0.05, marking the significance level for detecting deviations from the assumption of non-avoidance.

## References

Alberts, S. C. (2019). Social influences on survival and reproduction: Insights from a long-term study of wild baboons. Journal of Animal Ecology, 88(1), 47–66. 10.1111/1365-2656.12887

Allan, A. T. L., Bailey, A. L., & Hill, R. A. (2021). Consistency in the flight and visual orientation distances of habituated chacma baboons after an observed leopard predation. Do flight initiation distance methods always measure perceived predation risk? Ecology and Evolution, 11(21), 15404–15416. 10.1002/ece3.8237

Allan, A. T. L., LaBarge, L. R., Bailey, A. L., Jones, B., Mason, Z., Pinfield, T., Schröder, F., Whitaker, A., White, A. F., Wilkinson, H., & Hill, R. A. (2024). Behavioural compatibility, not fear, best predicts the looking patterns of chacma baboons. Communi-cations Biology, 7(1), 980. 10.1038/s42003-024-06657-w

Anderson, J. R. (1998). Sleep, sleeping sites, and sleep-related activities: Awakening to their significance. American Journal of Primatology, 46(1), 63–75. 10.1002/(SICI)1098-2345(1998)46:1<63::AID-AJP5>3.0.CO;2-T

Bailey, T. N. (1993). 9. Social Organization of Leopards. In The African Leopard (pp. 240–297). Columbia University Press. 10.7312/bail90198-010

Beaumont, R. A. R.□; P. B., & Bednarik, R. G. (2015). Concerning a cupule sequence on the edge of the Kalahari Desert in South Africa. 32(2), 163–177. 10.69978/rar.v32i2.136

Bedetti, A., Greyling, C., Paul, B., Blondeau, J., Clark, A., Malin, H., Horne, J., Makukule, R., Wilmot, J., Eggeling, T., Kern, J., & Henley, M. (2020). System for Elephant Ear-pattern Knowledge (SEEK) to identify individual African elephants. Pachyderm, 61, 63–77. 10.69649/pachyderm.v61i.65

Berger, J. (1979). “Predator Harassment” as a Defensive Strategy in Ungulates. American Midland Naturalist, 102(1), 197. 10.2307/2425087

Bersacola, E., Hill, C. M., Nijman, V., & Hockings, K. J. (2022). Examining primate community occurrence patterns in agroforest landscapes using arboreal and terrestrial camera traps. Landscape Ecology, 37(12), 3103–3121. 10.1007/s10980-022-01524-7

Bezerra, B. M., & Souto, A. (2008). Structure and Usage of the Vocal Repertoire of Callithrix jacchus. International Journal of Primatology, 29(3), 671–701. 10.1007/s10764-008-9250-0

Bidner, L. R., Matsumoto-Oda, A., & Isbell, L. A. (2018). The role of sleeping sites in the predator-prey dynamics of leopards and olive baboons. American Journal of Primatology, 80(12). 10.1002/ajp.22932

Blount, J. D., Chynoweth, M. W., Green, A. M., & Şekercioğlu, Ç. H. (2021). Review: COVID-19 highlights the importance of camera traps for wildlife conservation research and management. Biological Conservation, 256, 108984. 10.1016/j.biocon.2021.108984

Boyer-Ontl, K. M., & Pruetz, J. D. (2014). Giving the Forest Eyes: The Benefits of Using Camera Traps to Study Unhabituated Chimpanzees (*Pan troglodytes verus*) in Southeastern Senegal. International Journal of Primatology, 35(5), 881–894. 10.1007/s10764-014-9783-3

Broekhuis, F., Cozzi, G., Valeix, M., McNutt, J. W., & Macdonald, D. W. (2013). Risk avoidance in sympatric large carnivores: reactive or predictive? Journal of Animal Ecology, 82(5), 1098–1105. 10.1111/1365-2656.12077

Busse, C. D. (1980). Leopard and Lion predation upon Chacma Baboons living in the Moremi Wildlife Reserve. Botswana Society, 12, 15–21.

Chassagneux, A., Calenge, C., Marchand, P., Richard, E., Guillaumat, E., Baubet, E., & Saïd, S. (2020). Should I stay or should I go? Determinants of immediate and de-layed movement responses of female red deer (*Cervus elaphus*) to drive hunts. PLOS ONE, 15(3), e0228865. 10.1371/journal.pone.0228865

Courbin, N., Loveridge, A. J., Fritz, H., Macdonald, D. W., Patin, R., Valeix, M., & Chamaillé-Jammes, S. (2018). Zebra diel migrations reduce encounter risk with lions at night. Journal of Animal Ecology, 88(1), 92–101. 10.1111/1365-2656.12910

Courbin, N., Loveridge, A. J., Macdonald, D. W., Fritz, H., Valeix, M., Makuwe, E. T., & Chamaillé-Jammes, S. (2016). Reactive responses of zebras to lion encounters shape their predator–prey space game at large scale. Oikos, 125(6), 829–838. 10.1111/oik.02555

Cowlishaw, G. (1994). Vulnerability To Predation in Baboon Populations. Behaviour, 131(3–4), 293–304. 10.1163/156853994X00488

Cowlishaw, G. (1997). Trade-offs between foraging and predation risk determine habitat use in a desert baboon population. Animal Behaviour, 53(4), 667–686. 10.1006/anbe.1996.0298

Creel, S. (2018). The control of risk hypothesis: reactive vs. proactive antipredator responses and stress-mediated vs. food-mediated costs of response. Ecology Letters, 21(7), 947–956. 10.1111/ele.12975

Creel, S., & Christianson, D. (2008). Relationships between direct predation and risk effects. Trends in Ecology & Evolution, 23(4), 194–201. 10.1016/j.tree.2007.12.004

Creel, S., Schuette, P., & Christianson, D. (2014). Effects of predation risk on group size, vigilance, and foraging behavior in an African ungulate community. Behavioral Ecology, 25(4), 773–784. 10.1093/beheco/aru050

Creel, S., & Winnie, J. A. (2005). Responses of elk herd size to fine-scale spatial and temporal variation in the risk of predation by wolves. Animal Behaviour, 69(5), 1181–1189. 10.1016/j.anbehav.2004.07.022

Cusack, J. J., Dickman, A. J., Kalyahe, M., Rowcliffe, J. M., Carbone, C., MacDonald, D. W., & Coulson, T. (2017). Revealing kleptoparasitic and predatory tendencies in an African mammal community using camera traps: a comparison of spatiotemporal approaches. Oikos, 126(6), 812–822. 10.1111/oik.03403

Davis, A. L. V., Scholtz, C. H., Kryger, U., Deschodt, C. M., & Strümpher, W. P. (2010). Dung Beetle Assemblage Structure in Tswalu Kalahari Reserve: Responses to a Mosaic of Landscape Types, Vegetation Communities, and Dung Types. Environmental Entomology, 39(3), 811–820. 10.1603/EN09256

Dill, L. M., & Frid, A. (2020). Behaviourally mediated biases in transect surveys: a predation risk sensitivity approach. Canadian Journal of Zoology, 98(11), 697–704. 10.1139/cjz-2020-0039

Dröge, E., Creel, S., Becker, M. S., & M’soka, J. (2017). Spatial and temporal avoidance of risk within a large carnivore guild. Ecology and Evolution, 7(1), 189–199. 10.1002/ece3.2616

Druelle, F., Aerts, P., Ngawolo, J. C. B., & Narat, V. (2020). Impressive Arboreal Gap-Crossing Behaviors in Wild Bonobos, (Pan paniscus). International Journal of Primatology, 41(1), 129–140. 10.1007/s10764-020-00140-z

Edwards, S., Gange, A. C., & Wiesel, I. (2016). An oasis in the desert: The potential of water sources as camera trap sites in arid environments for surveying a carnivore guild. Journal of Arid Environments, 124, 304–309. 10.1016/j.jaridenv.2015.09.009

Elliot, N. B., Cushman, S. A., Loveridge, A. J., Mtare, G., & Macdonald, D. W. (2014). Movements vary according to dispersal stage, group size, and rainfall: the case of the African lion. Ecology, 95(10), 2860–2869. 10.1890/13-1793.1

Evans, B. E., Mosby, C. E., & Mortelliti, A. (2019). Assessing arrays of multiple trail cameras to detect North American mammals. PLOS ONE, 14(6), e0217543. 10.1371/journal.pone.0217543

Fehlmann, G., O’Riain, M. J., Kerr-Smith, C., Hailes, S., Luckman, A., Shepard, E. L. C., & King, A. J. (2017). Extreme behavioural shifts by baboons exploiting risky, resource-rich, human-modified environments. Scientific Reports, 7(1), 15057. 10.1038/s41598-017-14871-2

Fichtel, C., Perry, S., & Gros-Louis, J. (2005). Alarm calls of white-faced capuchin monkeys: an acoustic analysis. Animal Behaviour, 70(1), 165–176. 10.1016/j.anbehav.2004.09.020

Fischer, J., Metz, M., Cheney, D. L., & Seyfarth, R. M. (2001). Baboon responses to graded bark variants. Animal Behaviour, 61(5), 925–931. 10.1006/anbe.2000.1687

Foster, V. C., Sarmento, P., Sollmann, R., Tôrres, N., Jácomo, A. T. A., Negrões, N., Fonseca, C., & Silveira, L. (2013). Jaguar and Puma Activity Patterns and Predator-Prey Interactions in Four Brazilian Biomes. Biotropica, 45(3), 373–379. 10.1111/btp.12021

Galbany, J., Tung, J., Altmann, J., & Alberts, S. C. (2015). Canine Length in Wild Male Baboons: Maturation, Aging and Social Dominance Rank. PLOS ONE, 10(5), e0126415. 10.1371/journal.pone.0126415

Grolemund, G., & Wickham, H. (2011). Dates and Times Made Easy with lubridate. Journal of Statistical Software, 40(3). 10.18637/jss.v040.i03

Gursky, S. L. (2007). The Response of Spectral Tarsiers Toward Avian and Terrestrial Predators. In Primate Anti-Predator Strategies (pp. 241–252). Springer US. 10.1007/978-0-387-34810-0_11

Hammond, P., Gaynor, K., Easter, T., Biro, D., & Carvalho, S. (2025). Landscape-Scale Effects of Season and Predation Risk on the Terrestrial Behavior of Chacma Ba-boons (*Papio ursinus*). American Journal of Biological Anthropology, 186(4). 10.1002/ajpa.70052

Hammond, P., Lewis-Bevan, L., Biro, D., & Carvalho, S. (2022). Risk perception and terrestriality in primates: A quasi-experiment through habituation of chacma baboons (*Papio ursinus*) in Gorongosa National Park, Mozambique. American Journal of Bio-logical Anthropology, 179(1), 48–59. 10.1002/ajpa.24567

Harvell, C. D. (1990). The Ecology and Evolution of Inducible Defenses. The Quarterly Review of Biology, 65(3), 323–340. 10.1086/416841

Harvey, P. H., & Clutton-Brock, T. H. (1981). Primate home-range size and metabolic needs. Behavioral Ecology and Sociobiology, 8(2), 151–155. 10.1007/BF00300828

Hayward a, M. W., Henschel, P., O’Brien, J., Hofmeyr, M., Balme, G., & Kerley, G. I. H. (2006). Prey preferences of the leopard (*Panthera pardus*). Journal of Zoology, 270(2), 298–313. 10.1111/j.1469-7998.2006.00139.x

Hayward b, M. W., Hofmeyr, M., O’Brien, J., & Kerley, G. I. H. (2006). Prey preferences of the cheetah (*Acinonyx jubatus*) (Felidae: Carnivora): morphological limitations or the need to capture rapidly consumable prey before kleptoparasites arrive? Journal of Zoology, 270(4), 615–627. 10.1111/j.1469-7998.2006.00184.x

Hayward c, M. W., O’Brien, J., Hofmeyr, M., & Kerley, G. I. H. (2006). Prey Preferences of the African wild dog (*Lycaon pictus*) Canidae Canidae: Carnivora: Ecological requirments for conservation. Journal of Mammalogy, 87(6), 1122–1131. 10.1644/05-MAMM-A-304R2.1

Herrera, D. J., Moore, S. M., Herrmann, V., McShea, W. J., & Cove, M. V. (2021). A shot in the dark: White and infrared LED flash camera traps yield similar detection probabilities for common urban mammal species. Hystrix, the Italian Journal of Mammalogy, 32(1), 72–75. 10.4404/hystrix-00429-2021

Hill, R. A., & Weingrill, T. (2007). Predation Risk and Habitat Use in Chacma Baboons (*Papio hamadryas ursinus*). In Primate Anti-Predator Strategies (pp. 339–354). Springer US. 10.1007/978-0-387-34810-0_16

Hoffman, T. S., & O’riain, M. Justin. (2012). Troop Size and Human-Modified Habitat Affect the Ranging Patterns of a Chacma Baboon Population in the Cape Peninsula, South Africa. American Journal of Primatology, 74(9), 853–863. 10.1002/ajp.22040

Horrocks, J. A., & Hunte, W. (1986). Sentinel behaviour in vervet monkeys: Who sees whom first? Animal Behaviour, 34(5), 1566–1568. 10.1016/S0003-3472(86)80226-3

Hubel, T. Y., Golabek, K. A., Rafiq, K., McNutt, J. W., & Wilson, A. M. (2018). Movement patterns and athletic performance of leopards in the Okavango Delta. Proceedings of the Royal Society B: Biological Sciences, 285(1877), 20172622. 10.1098/rspb.2017.2622

Humphries, D. A., & Driver, P. M. (1967). Erratic Display as a Device against Predators. Science, 156(3783), 1767–1768. 10.1126/science.156.3783.1767

Isbell, L. A., Bidner, L. R., Van Cleave, E. K., Matsumoto-Oda, A., & Crofoot, M. C. (2018). GPS-identified vulnerabilities of savannah-woodland primates to leopard predation and their implications for early hominins. Journal of Human Evolution, 118, 1–13. 10.1016/j.jhevol.2018.02.003

James, D., & Hornik, K. (1999). chron: Chronological Objects which Can Handle Dates and Times. In CRAN: Contributed Packages. 10.32614/CRAN.package.chron

Johnson, S. E. (2003). Life history and the competitive environment: trajectories of growth, maturation, and reproductive output among Chacma baboons. American Journal of Physical Anthropology, 120(1), 83–98. 10.1002/ajpa.10139

Jooste, E., Pitman, R. T., van Hoven, W., & Swanepoel, L. H. (2013). Unusually High Predation on Chacma Baboons (*Papio ursinus*) by Female Leopards (*Panthera pardus*) in the Waterberg Mountains, South Africa. Folia Primatologica, 83(3–6), 353–360. 10.1159/000339644

Kappeler, P. M., & Van Schaik, C. P. (2002). Evolution of Primate Social Systems. In International Journal of Primatology (Vol. 23, Number 4).

Keim, J. L., Lele, S. R., DeWitt, P. D., Fitzpatrick, J. J., & Jenni, N. S. (2019). Estimating the intensity of use by interacting predators and prey using camera traps. Journal of Animal Ecology, 88(5), 690–701. 10.1111/1365-2656.12960

Kerbs, John. R., & Davies, Nicholas. B. (2009). Behavioural ecology: an evolutionary approach. John Wiley & Sons.

Kie, J. G. (1999). Optimal Foraging and Risk of Predation: Effects on Behavior and Social Structure in Ungulates. Journal of Mammalogy, 80(4), 1114–1129. 10.2307/1383163

Kolowski, J. M., Katan, D., Theis, K. R., & Holekamp, K. E. (2007). Daily Patterns of Activity in the Spotted Hyena. Journal of Mammalogy, 88(4), 1017–1028. 10.1644/06-MAMM-A-143R.1

Kramer, D. L., & Bonenfant, M. (1997). Direction of predator approach and the decision to flee to a refuge. Animal Behaviour, 54(2), 289–295. 10.1006/anbe.1996.0360

LaBarge, L. R., Allan, A. T. L., Berman, C. M., Hill, R. A., & Margulis, S. W. (2022). Cortisol metabolites vary with environmental conditions, predation risk, and human shields in a wild primate, (*Cercopithecus albogularis*). Hormones and Behavior, 145, 105237. 10.1016/j.yhbeh.2022.105237

Laundré, J. W., Hernández, L., & Altendorf, K. B. (2001). Wolves, elk, and bison: reestablishing the “landscape of fear” in Yellowstone National Park, U.S.A. Canadian Journal of Zoology, 79(8), 1401–1409. 10.1139/z01-094

Li, Y. (2007). Terrestriality and tree stratum use in a group of Sichuan snub-nosed monkeys. Primates, 48(3), 197–207. 10.1007/s10329-006-0035-9

Lima, S. L., & Dill, L. M. (1990). Behavioral decisions made under the risk of predation: a review and prospectus. Canadian Journal of Zoology, 68(4), 619–640. 10.1139/z90-092

Linkie, M., & Ridout, M. S. (2011). Assessing tiger–prey interactions in Sumatran rainforests. Journal of Zoology, 284(3), 224–229. 10.1111/j.1469-7998.2011.00801.x

MacCormick, H. A., MacNulty, D. R., Bosacker, A. L., Lehman, C., Bailey, A., Anthony Collins, D., & Packer, C. (2012). Male and female aggression: lessons from sex, rank, age, and injury in olive baboons. Behavioral Ecology, 23(3), 684–691. 10.1093/beheco/ars021

Martins, Q., & Harris, S. (2013). Movement, activity and hunting behaviour of leopards in the Cederberg mountains, South Africa. African Journal of Ecology, 51(4), 571–579. 10.1111/aje.12068

Meek, P. D., Ballard, G., Claridge, A., Kays, R., Moseby, K., O’Brien, T., O’Connell, A., Sanderson, J., Swann, D. E., Tobler, M., & Townsend, S. (2014). Recommended guiding principles for reporting on camera trapping research. Biodiversity and Conservation, 23(9), 2321–2343. 10.1007/s10531-014-0712-8

Meek, P. D., Ballard, G., Fleming, P., & Falzon, G. (2016). Are we getting the full picture? Animal responses to camera traps and implications for predator studies. Ecology and Evolution, 6(10), 3216–3225. 10.1002/ece3.2111

Musyoki, C. M., & Strum, S. C. (2016). Spatial and temporal patterns of home range use by olive baboons (*Papio anubis*) in eastern Laikipia, Kenya. African Journal of Ecology, 54(3), 349–356. 10.1111/aje.12294

Newey, S., Davidson, P., Nazir, S., Fairhurst, G., Verdicchio, F., Irvine, R. J., & van der Wal, R. (2015). Limitations of recreational camera traps for wildlife management and conservation research: A practitioner’s perspective. Ambio, 44, 624–635. 10.1007/s13280-015-0713-1

Niedballa, J., Wilting, A., Sollmann, R., Hofer, H., & Courtiol, A. (2019). Assessing analytical methods for detecting spatiotemporal interactions between species from camera trapping data. Remote Sensing in Ecology and Conservation, 5(3), 272–285. 10.1002/rse2.107

Noser, R., & Byrne, R. W. (2010). How do wild baboons (*Papio ursinus*) plan their routes? Travel among multiple high-quality food sources with inter-group competition. Animal Cognition, 13(1), 145–155. 10.1007/s10071-009-0254-8

Nowak, K., le Roux, A., Richards, S. A., Scheijen, C. P. J., & Hill, R. A. (2014). Human observers impact habituated samango monkeys’ perceived landscape of fear. Behavioral Ecology, 25(5), 1199–1204. 10.1093/beheco/aru110

Palmer, M. S., Gaynor, K. M., Becker, J. A., Abraham, J. O., Mumma, M. A., & Pringle, R. M. (2022). Dynamic landscapes of fear: understanding spatiotemporal risk. Trends in Ecology & Evolution, 37(10), 911–925. 10.1016/j.tree.2022.06.007

Palmer, M. S., & Packer, C. (2021). Reactive anti-predator behavioral strategy shaped by predator characteristics. PLOS ONE, 16(8), e0256147. 10.1371/journal.pone.0256147

Pardo, L. E., Bombaci, S., Huebner, S. E., Somers, M. J., Fritz, H., Downs, C., Guthmann, A., Hetem, R. S., Keith, M., le Roux, A., Mgqatsa, N., Packer, C., Palmer, M. S., Parker, D. M., Peel, M., Slotow, R., Maartin Strauss, W., Swanepoel, L., Tambling, C., … Venter, J. A. (2021). Snapshot Safari: A large-scale collaborative to monitor Africa’s remarkable biodiversity. South African Journal of Science, 117(1–2). 10.17159/SAJS.2021/8134

Rutherford, M. C., Mucina, L., Lötter, M. C., Bredenkamp, G. J., Smit, J. H. L., Scott-Shaw, C. R., Hoare, D. B., Goodman, P. S., Bezuidenhout, H., Scott, L., Ellis, F., Powrie, L., Siebert, F., Mostert, T. H., Henning, B. J., Ventner, C. E., Camp, K. G. T., Siebert, S., Matthews, W., & Hurter, J. (2006). Savanna Biome. In L. Mucina & M. C. Rutherford (Eds.), The Vegetation of South Africa, Lesotho and Swaziland. SANBI.

Sankaran, M., Hanan, N. P., Scholes, R. J., Ratnam, J., Augustine, D. J., Cade, B. S., Gignoux, J., Higgins, S. I., Le Roux, X., Ludwig, F., Ardo, J., Banyikwa, F., Bronn, A., Bucini, G., Caylor, K. K., Coughenour, M. B., Diouf, A., Ekaya, W., Feral, C. J., … Zambatis, N. (2005). Determinants of woody cover in African savannas. Nature, 438(7069), 846–849. 10.1038/nature04070

Say-Sallaz, E., Chamaillé-Jammes, S., Périquet, S., Loveridge, A. J., Macdonald, D. W., Antonio, A., Fritz, H., & Valeix, M. (2023). Large carnivore dangerousness affects the reactive spatial response of prey. Animal Behaviour, 202, 149–162. 10.1016/j.anbehav.2023.05.014

Scholes, R. J., & Archer, S. R. (1997). Tree-Grass Interactions in Savannas. Annual Review of Ecology and Systematics, 28(1), 517–544. 10.1146/annurev.ecolsys.28.1.517

Schreier, A. L., & Swedell, L. (2012). Ecology and sociality in a multilevel society: Ecological determinants of spatial cohesion in hamadryas baboons. American Journal of Physical Anthropology, 148(4), 580–588. 10.1002/ajpa.22076

Searle, C. E., Smit, J. B., Cusack, J. J., Strampelli, P., Grau, A., Mkuburo, L., Macdonald, D. W., Loveridge, A. J., & Dickman, A. J. (2021). Temporal partitioning and spatio-temporal avoidance among large carnivores in a human-impacted African landscape. PLOS ONE, 16(9), e0256876. 10.1371/journal.pone.0256876

Seyfarth, R. M., Cheney, D. L., & Marler, P. (1980). Vervet monkey alarm calls: Semantic communication in a free-ranging primate. Animal Behaviour, 28(4), 1070–1094. 10.1016/S0003-3472(80)80097-2

Shannon, G., Cordes, L. S., Hardy, A. R., Angeloni, L. M., & Crooks, K. R. (2014). Behavioral Responses Associated with a Human-Mediated Predator Shelter. PLoS ONE, 9(4), e94630. 10.1371/journal.pone.0094630

Si, X., Kays, R., & Ding, P. (2014). How long is enough to detect terrestrial animals? Estimating the minimum trapping effort on camera traps. PeerJ, 2, e374. 10.7717/peerj.374

Slater, K., Barrett, A., & Brown, L. R. (2018). Home range utilization by chacma baboon (Papio ursinus) troops on Suikerbosrand Nature Reserve, South Africa. PLOS ONE, 13(3), e0194717. 10.1371/journal.pone.0194717

Smith, J. A., Suraci, J. P., Hunter, J. S., Gaynor, K. M., Keller, C. B., Palmer, M. S., Atkins, J. L., Castañeda, I., Cherry, M. J., Garvey, P. M., Huebner, S. E., Morin, D. J., Teckentrup, L., Weterings, M. J. A., & Beaudrot, L. (2020). Zooming in on mechanistic predator–prey ecology: Integrating camera traps with experimental methods to re-veal the drivers of ecological interactions. Journal of Animal Ecology, 89(9), 1997–2012. 10.1111/1365-2656.13264

Stone, O. M. L., Laffan, S. W., Curnoe, D., & Herries, A. I. R. (2013). The Spatial Distribution of Chacma Baboon (Papio ursinus) Habitat Based on an Environmental Envelope Model. International Journal of Primatology, 34(2), 407–422. 10.1007/s10764-013-9669-9

Sugardjito, J. (1983). Selecting nest-sites of sumatran orangutans, (*Pongo pygmaeus abelii*) in the Gunung Leuser National Park, Indonesia. Primates, 24(4), 467–474. 10.1007/BF02381680

Suire, A., Kunita, I., Harel, R., Crofoot, M., Mutinda, M., Kamau, M., Hassel, J. M., Murray, S., Kawamura, S., & Matsumoto-Oda, A. (2023). Estimating individual exposure to predation risk in group-living baboons, Papio anubis. PLOS ONE, 18(11), e0287357. 10.1371/journal.pone.0287357

Suscke, P., Presotto, A., & Izar, P. (2021). The role of hunting on *Sapajus xanthosternos’’landscape of fear in the Atlantic Forest, Brazil*. American Journal of Primatology, 83(5). 10.1002/ajp.23243

Swanson, A., Kosmala, M., Lintott, C., Simpson, R., Smith, A., & Packer, C. (2015). Snapshot Serengeti, high-frequency annotated camera trap images of 40 mammalian species in an African savanna. Scientific Data, 2(1), 150026. 10.1038/sdata.2015.26

Swinkels, C., van der Wal, J. E. M., Stinn, C., Monteza-Moreno, C. M., & Jansen, P. A. (2023). Prey tracking and predator avoidance in a Neotropical moist forest: a cam-era-trapping approach. Journal of Mammalogy, 104(1), 137–145. 10.1093/jmammal/gyac091

Szott, I. D., Pretorius, Y., Ganswindt, A., & Koyama, N. F. (2020). Physiological stress response of African elephants to wildlife tourism in Madikwe Game Reserve, South Africa. Wildlife Research, 47(1), 34. 10.1071/WR19045

Thaker, M., Vanak, A. T., Owen, C. R., Ogden, M. B., Niemann, S. M., & Slotow, R. (2011). Minimizing predation risk in a landscape of multiple predators: Effects on the spatial distribution of African ungulates. Ecology, 92(2), 398–407. 10.1890/10-0126.1

Tjaden-McClement, K., Gharajehdaghipour, T., Shores, C., White, S., Steenweg, R., Bourbonnais, M., Konanz, Z., & Burton, A. C. (2025). Mixed evidence for disturbance-mediated apparent competition for declining caribou in western British Columbia, Canada. The Journal of Wildlife Management, 89(6). 10.1002/jwmg.70040

Trinkel, M. (2010). Prey selection and prey preferences of spotted hyenas (*Crocuta crocuta*) in the Etosha National Park, Namibia. Ecological Research, 25(2), 413–417. 10.1007/s11284-009-0669-3

van Schaik, C. P., Bshary, R., Wagner, G., & Cunha, F. (2022). Male anti-predation services in primates as costly signalling? A comparative analysis and review. Ethology, 128(1), 1–14. 10.1111/eth.13233

Virgadamo, P., Hodosh, M., Povar, M., & Shklar, G. (1972). The Dentition of Papio anubis. Journal of Dental Research, 51(5), 1338–1345. 10.1177/00220345720510051501

Von Hippel, F. A. (1998). Use of sleeping trees by black and white Colobus monkeys (Colobus guereza) in the Kakamega Forest, Kenya. American Journal of Primatology, 45(3), 281–290. 10.1002/(SICI)1098-2345(1998)45:3<281::AID-AJP4>3.0.CO;2-S

Wentworth, J. C., Tambling, C. J., & Kerley, G. I. H. (2011). Evidence for prey selection by spotted hyaena in the Eastern Cape, South Africa. Acta Theriologica, 56(4), 389–392. 10.1007/s13364-011-0033-1

Weterings, M. J. A., Zaccaroni, M., van der Koore, N., Zijlstra, L. M., Kuipers, H. J., van Langevelde, F., & van Wieren, S. E. (2016). Strong reactive movement response of the medium-sized European hare to elevated predation risk in short vegetation. Animal Behaviour, 115, 107–114. 10.1016/j.anbehav.2016.03.011

Wickham, H. (2009). stringr: Simple, Consistent Wrappers for Common String Operations. In CRAN: Contributed Packages. 10.32614/CRAN.package.stringr

Wickham, H., Vaughan, D., & Girlich, M. (2014). tidyr: Tidy Messy Data. In CRAN: Contributed Packages. 10.32614/CRAN.package.tidyr

Willems, E. P., & Hill, R. A. (2009). Predator-specific landscapes of fear and resource distribution: effects on spatial range use. Ecology, 90(2), 546–555. 10.1890/08-0765.1

Zhang, Y., & Zhao, Q. (2023). What is a Randomization Test? Journal of the American Statistical Association, 118(544), 2928–2942. 10.1080/01621459.2023.2199814

Zinner, D., Buba, U., Nash, S., & Roos, C. (2011). Pan-African Voyagers: The Phy-logeography of Baboons. In Primates of Gashaka (pp. 319–358). Springer New York. 10.1007/978-1-4419-7403-7_7

